# Analyzing photoactivation with diffusion models to study transport in the Endoplasmic Reticulum network

**DOI:** 10.1101/2023.11.14.567043

**Authors:** M. Dora, F. Paquin-Lefebvre, D. Holcman

## Abstract

Photoactivation is a paradigm consisting in local molecular fluorescent activation by laser illumination in a chosen region (source) while measuring the concentration at a target region. Data-driven modeling is concern with the following questions: how from the measurement in these two regions, is it possible to infer the properties of molecular propagation? How is it possible to use such reponses to infer motions occurring in networks such as the endoplasmic reticulum? In this article, we present a data-driven analysis based on diffusion-transport models and numerical simulations to interpret the photoactivation dynamics and extract biophysical parameters. Finally we discuss modeling approaches to reconstruct local network properties from photoactivation transients.

## 1 Introduction

How molecules, ions or proteins spread across cell compartments remains an active field of cell biology. Molecules can move passively as Brownian motion but can also be transported actively by molecular motors from one region to another. Yet exploring the exact propagation mode is not straightforward as it requires combining molecular labeling with refined live super-resolution microscopy.

More than 20 year ago was developed a key approach called fluorescence recovery after photobleaching (FRAP), which consists in photo-bleaching a region labeled with a fluorescent marker and observing how this region gets re-populated [1]. This approach in combination with modeling propagation as diffusion provided a first tool to fit experimental recovery curves by a diffusion model, allowing estimating an effective diffusion coefficient and the fraction of immobile molecules. Later on, an opposite paradigm emerged [2, 3] because of a new photoactivable (PA) molecule PA-GFP (Green Fluorescent Protein), where a population was photoactivated. For that case experimental data is obtained by counting the arrival of activated molecules in a given region. Photoactivation consists in locally activating fluorescent proteins by laser illumination in a chosen region (source) and measuring the propagation at a target region located at a distance apart from the source [4], thus providing information about first arrival times and also biophysical parameters such as diffusion.

While these experiments are not too difficult to perform in the cytosol of a cell, they remain quite tedious in organelles such as the endoplasmic reticulum which consists of a thin network being constantly reshaped, with tubule size lying under the diffraction limit. Analyzing fluorescence data collected from population photoactivation in the ER remains challenging due to several difficulties associated with noisy signals, segmentation of the ER tubules and the lack of a defined biophysical model to describe lumen and membrane motion. Various imaging approaches have been developed to denoise these images: first is the variance stabilizing transformation followed by additive white noise removal [5], algorithms specifically designed for Poisson-Gaussian mixtures [6], and deep learning models based on convolution neural networks [7, 8]. These approaches are not the subject of the current manuscript and will be reviewed somewhere else.

However, the experimental approaches mentioned above are in contrast with single particle trajectories (SPTs) where each molecule is activated at a random time and at a random position, leading to a random sample of trajectories. Interestingly, previous work with SPTs reveal active motion in the ER tubules, while passive diffusion in nodes. But today some discrepancies persist in the description of motion between the population versus the individual trajectory approaches [9, 10].

In this manuscript, we shall present modeling approaches used to estimate and to interpret transient rates at which molecules disappear from one region to appear in another. We shall further discuss how propagation in the endoplasmic reticulum can be studied by solving simple planar diffusion models. We will also show how network simulations can be employed to explore the consequence of an active flow between ER nodes. Lastly we will discuss simulations to extract arrival times to ER nodes.

## 2 Principles of photoactivation and source-target interaction

Photoactivation consists in locally activating fluorescent proteins by laser illumination in a chosen region (source)(Fig. 1A) and measuring the propagation at a target region located at a distance *d* apart from the source. The source and the target are disks of small radius (few microns), that can be adjusted from the microscope aperture even if located on a network such as the ER (Fig. 1B-C). This procedure allows recording the fluorescence signal *f* (*d, t*) on the target at time t. The transient fluorescence time course carries biophysical information, such as the propagation mode (diffusion, drift, active motion or any combination) and the local structure of the underlying network with possible inhomogeneous node-tubule distribution, that should be extracted from the function *f* (*d, t*). However, this procedure requires a model for *f*, in addition to a statistical method to extract the relevant parameters.

**Figure 1:**
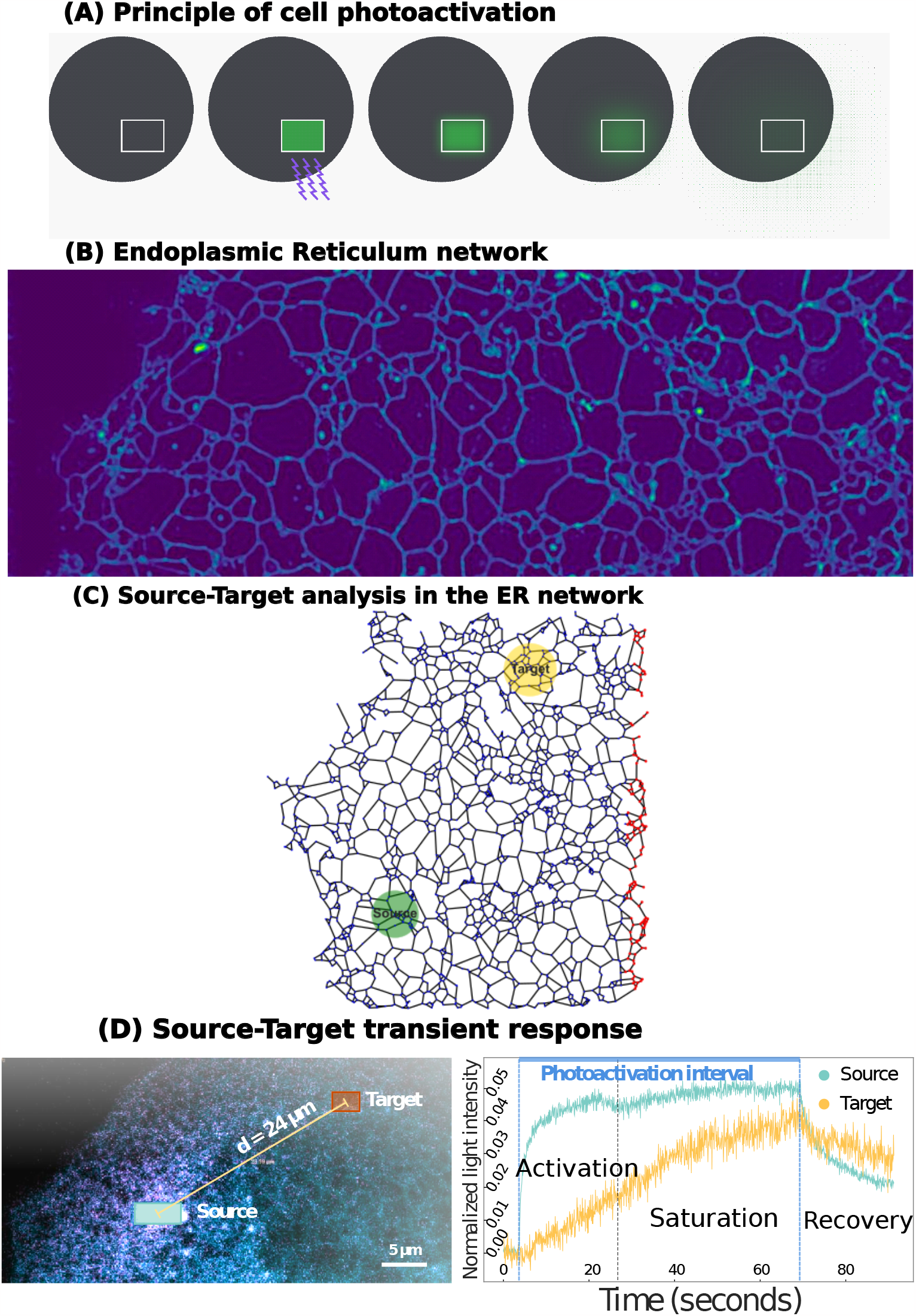
Principle of photoactivation in the ER. **A**. A subregion of a cell (round scheme) is activated by a laser, followed by the spread of activated particles in the cell and their disappearance from the source region. **B**. Example of an Endoplasmic Reticulum network. **C**. Principle of source-target photoactivation analysis: a source region is photoactivated while the fluorescent signal is analyzed at the target region. **D**. Transient responses of photoactivation at the source (blue) and target (yellow). The dynamics is decomposed into three phases: 1-Transient activation starts when fluorescent molecules are activated, until 2-A transient steady-state is achieved (saturation). 3-The last phase starts when the laser stops, leading to a different decay in both regions.

Photoactivation can be decomposed into three phases. There is first a fast rising phase where dye molecules are photoactivated (Fig. 1D), during which some molecules can be exchanged with the neighboring environment and which lasts until a saturation is reached (plateau in Fig. 1D). In the second phase, the signal saturates until the laser is stopped. During these two phases, the signal increases at the target, interestingly sometimes close to the saturation amplitude of the source.

The third phase starts when the laser excitation stops, it is characterized by a relaxation of the activated molecule toward the rest of the ER, leading to a decay of the concentration as the activated molecules spread inside the network. The three phases can be recorded at any distance from the source (from few to tens of microns or more). The goal of a modeling approach is to extract from the ensemble of response the type of motion occurring in the ER. The area of the source and target can be chosen containing one or multiple ER-nodes. For a size of several microns, the molecule can be locally mixed and it would then be possible to use a continuous approximation of the ER geometry such as the two-dimensional plane. Another possibility is to simulate the ER as a network where the mean number of nodes (degree) is equal to three [9]. Photo-activation modeling can thus be used to quantify transport within the ER. In brief, we propose to study the local (few microns) and far away transport (tens of micron or more). We summarize below the possible local transport hypotheses inside the ER.

### 2.1 Molecular transport hypotheses in the ER

We summarize here various modes of transport that could occur in the ER, which can be explored through modeling and simulations.

1. **Large scale diffusion**. Molecular transport could be dominated by diffusion at both small and large scales. In that case, computing a single diffusion coefficient is sufficient to characterize the entire dynamics. This hypothesis was initially tested using FRAP and lead to a diffusion coefficient of *D* = 1.65 *µ*m^2^*/*s [11].
2. **Spatial dependent diffusion**. Another possibility is a dynamics characterize by a spatial dependent diffusion, with the diffusion coefficient *D*(***x***) depending on the position ***x*** (that could represent the center of a disk of radius *a*, where *a* characterizing the resolution). The output of this analysis is a diffusion map [12] and in that case the diffusion coefficient *D*(***x***) characterizes the local transport. Spatial variation of the diffusivity should be explained: nodes could be of different sizes or contain fewer passages, or there could be changes in the density of nodes.
3. **Active drift and diffusion**. An active drift potentially depending on ATP could also be possible. This was observed in tubules using SPTs [9], but was not found at higher spatial scales.
4. **Alternate active drift in tubules** . If tubules can let molecules move in only one direction at a time by a yet unknown mechanism, then it is possible to anticipate a motion where molecules move alternativaly and this would lead to molecular packets.
5. **ER membrane transport**. Transport on ER membrane could be different from the lumen motion [9].
6. **Asymmetry transport**. Some asymmetry could exist between diffusion or active transport at region near the nucleus vs cell periphery.

We will show below how photoactivation (PA) can serve to test the different transport hypotheses.

### 2.2 Fluorescent Mean Square Displacement analysis

In this section, we recall diffusion models that are used to characterize the type of molecular motion in the ER lumen. We further mention how parameters should be computed to avoid certain pitfalls.

To quantify the nature of the motion from the PA data, one method consists in fitting for each cell the effective Mean Square Displacement (MSD) curve describing the time evolution of the average distance travelled by a protein released from the PA source (Fig. 2A-D). Is it worth stating the assumption about local isotropy that was tested: for collecting data, we use PA responses in *p* different small regions located at a distance *r* from the source and the average fluoresence *r*^2^*u*_*i*_(*r, t*) was recorded for *i* = 1, …, *p*. After varying the distance *r*, a collection of points is obtained (Fig. 2D).

**Figure 2:**
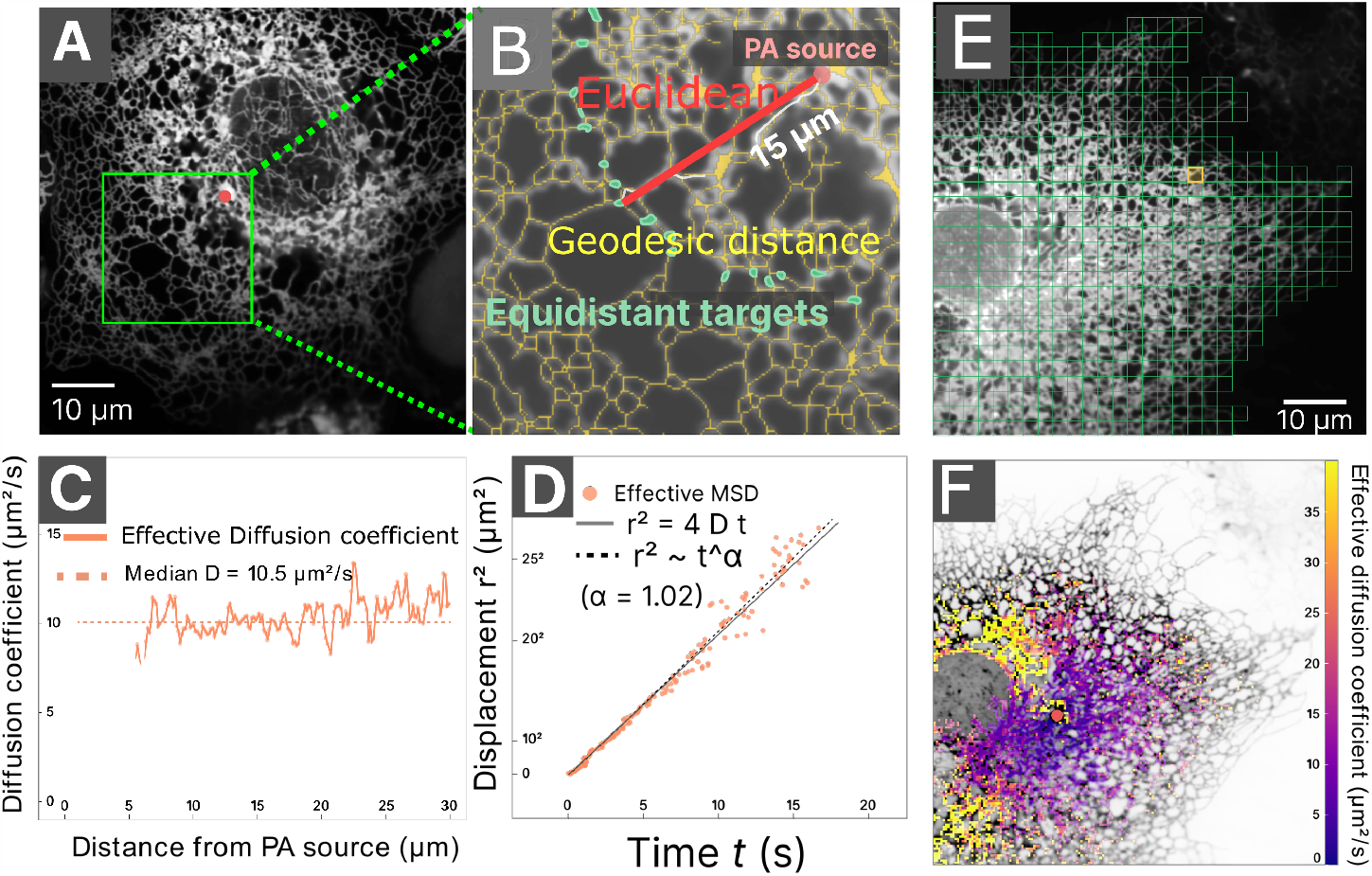
Diffusion analysis of large scale ER luminal dynamics. **A-B** ER structure of a COS-7 cell showing the location of photoactivation (red dot). ER subdomains (green regions) all equidistant from the photoactivation region (the distance calculated on the underlying ER morphology is *d* = 15 *µ*m). Here the Euclidean distance from the photoactivation region is 15 *µm* (red line). **C** Effective diffusion coefficient vs the distance *d* from the photoactivation region, obtained from separate fits at each distance, for the model accounting for the ER morphology (orange) and neglecting it (light blue). The error bars represent the approximated confidence interval estimated by observed Fisher information (± 5 standard deviations). **D** Effective mean-squared displacement (MSD) obtained by ensemble estimate. **E** Subdivision of the ER with a grid of square bins. **F** Spatial map describing the effective diffusion coefficient computed from the mean arrival time of HaloTag-KDEL. The red dot indicates the PA source. The grid bin size is 400 nm (3 by 3 camera pixels), adpated from [10].

To evaluate possible deviations from classical diffusion, a fit of thee MSD point distribution curves is used with the function *ct*^*α*^ (Fig. 2D), where *α* is called the anomalous exponent. The exponent *α* = 1 is associated with classical diffusion, while it represents subdiffusion for *α <* 1 and superdiffusion for *α >* 1. This procedure can be repeated for each cell to determine the MSD exponent *α*.

It is interesting to note that how distances are measured does influence fitting procedure results. A first choice is the Euclidean distance, given by the shortest segment from the source to the target (red line in Fig. 2A1). Another possibility is to instead consider the geodesic distance, which corresponds to the length of the minimal path within the ER network (thus accounting for its morphology) connecting the source to the target (white path in Fig. 2A1). The geodesic distance is longer than the Euclidean distance.

For instance, by calculating the MSD curve with the geodesic distance, population statistics of cells expressing HaloTag-KDEL had an average MSD exponent of *α≈* 1, thus suggesting a description of luminal dynamics by classical diffusion over this spatiotemporal scale (Fig. 2E, green). However, neglecting the ER morphology and using the Euclidean distance resulted in an apparent super-diffusive behaviour with *α* ≈ 1.2 [10]. This discrepancy could create a confusion, arguing for a possible drift toward the peripheral ER [13, 14, 15, 16].

## 3 Predicting the rising phase of the target from source fitting within the diffusion regime

We present in this section various models that can be used to extract parameters from the early phases of photoactivation between two regions called the source (laser target) and a target region (a region located at a certain distance away).

In a first approximation, diffusion models have been used to extract the dynamics occurring in the ER lumen or membrane. A first consists replacing the complex ER network geometry by a two-dimensional planar geometry, thereby neglecting the local topology (three-way junctions) dynamics of tubules and the possible heterogeneity of ER distribution [10], or the anomalous restricted motion of junctions [17]. For example, the ER structure in COS-7 cells is sufficiently flat to be considered as planar [18].

However two-dimensional geometries are insufficient for three-dimensional ER networks, for which more appropriate approximations could be obtained by considering a fractal description with an average dimension *α* = 1.6 [17]. This representation accounts for the number *n*(*r*) of ER networks located in a ball of radius r, changing *n*(*r*) *≈ r*^*α*^ as *r* increases [17]. This representation results in an overall anomalous diffusion [19].

### 3.1 Diffusion model of photoactivation at the source

We first describe in this subsection a diffusion method to extract the effective diffusion coefficient using an approximation of the ER-network as a two-dimensional plane. In that case the structure of each node/sheet/tubule is not accounted for and simply replaced by a continuum of distances. Also, the distance between the source and the target is computed using the Euclidean norm, an over simplification as ER lengths could be longer and should thus be computed over network paths.

A diffusion model allows to predict the concentration changes at the target site (Fig. 2B) from the dynamics of the source. The fluorescence curve *ϕ*(*t*) at the source *S* of the photoactivation region of interest (ROI) is approximated as a sum of exponentially saturating functions

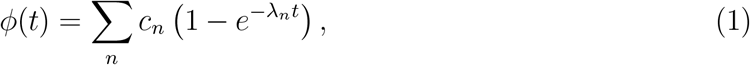

where the parameters (*c*_*n*_, *λ*_*n*_) are fitted to the empirical fluorescence (Fig. 2C). In practice a sum of 3 exponentials is sufficient. The next step is to use the diffusion equation in the plane to compute the concentration at the target (Fig. 2B). An optimal fit of the data allows to estimate the diffusion coefficient *D*. This approach considers that the recorded luminosity is proportional to the density of photoactivated particles.

### 3.2 Diffusion model of photoactivation at the target

The density of photoactivated particles at any point in space is obtained by solving the diffusion equation

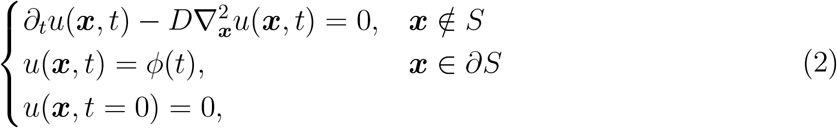

where the source effect *ϕ*(*t*) is imposed at the boundary of the source region *S*, with a zero initial condition. The solution can be expressed using a simpler auxiliary problem with a time independent boundary condition (called Duhamel’s principle [20])

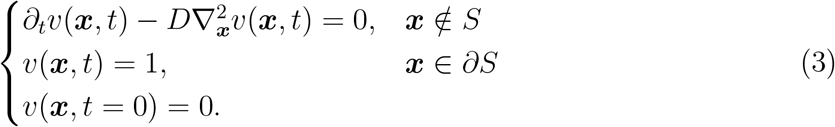

The solution of equation 2 is the convolution between the auxiliary solution *v*(*r, t*) and the time-varying boundary term:

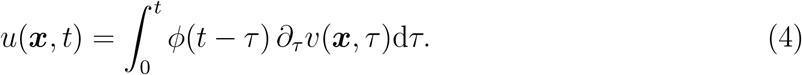

We now solve problem (3) in 2D with radial symmetry, where the source *S* consists of a disk of radius *a* (thus the domain is *r > a*) using the Laplace’s transform ℒ*{·}* . Taking then the Laplace transform [21] on both sides of (3) yields the modified Bessel equation of order 0,

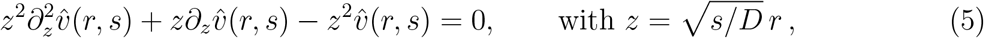

where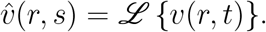. The solution is

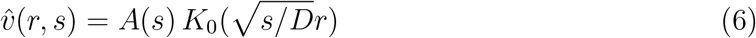

where *K*_0_(*z*) is the modified Bessel function of the second kind and order 0, while *A*(*s*) is a coefficient determined by imposing the boundary condition on *r* = *a*, which yields

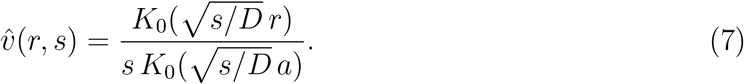

Finally the solution of the time-dependent problem is

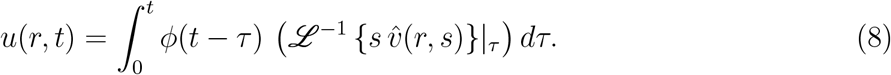

Eq. 8 is evaluated numerically, thus allowing to fit the photoactivation ROI at the target and recover an approximated diffusion coefficient *D*, which can be computed over the entire ER, leading to a diffusion map representation (Fig. 2E-F).

### 3.3 Numerical simulations of photoactivation with diffusion: exploring various target local geometries

To better characterize the photoactivation protocol various scenarios, each associated with a different target geometry, can be explored: 1-an open local domain 2-an absorbing domain 3-a partial absorbing domain (Fig. 4). Molecules are moving by diffusion inside a domain consisting of a large disk *B*(*O, R*) of radius *R* centered at the origin *O*, in which the source photoactivation region corresponds to the smaller disk *B*(*O, a*) of radius *a « R* (Fig. 4). Inside the source, molecules are activated by a laser during the time interval [0, *t*_*s*_], which can be modeled by a step function. The concentration of activated molecules is then measured over another disk *B*(*P, a*) located at a distance *L* = |*OP* | away from the source.

#### 3.3.1 Photoactivation diffusion model

We now present the diffusion equations that serve as a model of photoactivation. The concentration *u*(***x***, *t*) is modeled by the diffusion equation [22]

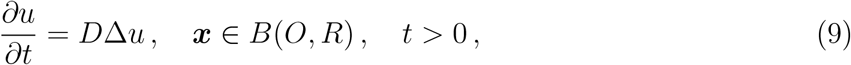

with reflective boundary conditions (no material escapes the domain)

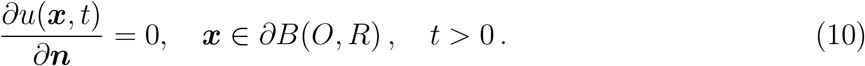

The laser source is activated during the time interval [0, *t*_*s*_], modeled by

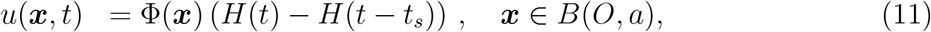

where the profile Φ (***x***) can be uniform or Gaussian. Here *H*(*t*) is the Heaviside function (*H*(*t*) = 0 for *t <* 0 and *H*(*t*) = 1 for *t >* 0). A modified diffusion equation has to be solve after a time *t > t*_*s*_, with *u*(***x***, *t*_*s*_) as an initial condition for again equation 9. However this may not be straightforward, and one alternative is to replace equations (9)-(11) by a single equation with an appropriate spatiotemporal source term:

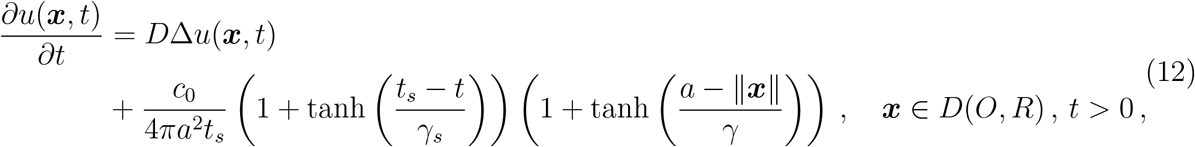

with zero initial condition *u*(***x***, *t*) = 0 and reflective boundary condition. The source term acts as a step function from time 0*≤ t≤ t*_*s*_ on the narrow disk Ω_*s*_ =*{* ***x***| 0*≤* || ***x*** ||*≤ a}* . The concentration at the target is computed by solving equation 12 and by computing the local concentration at the target *B*(*P, a*):

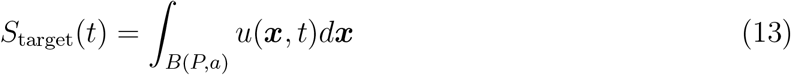

Similarly, at the source the total fluorescence is

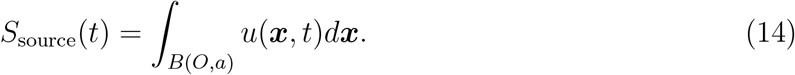

The source-target photo-activation responses are evaluated for three different cases:

1. **No further boundary conditions are imposed at the target site** *B*(*P, a*). Molecules can freely diffuse in and out of the target region.
2. **The target boundary consists of a dominant reflective part and a small freely diffusing one**. The target boundary decomposes into two pieces as 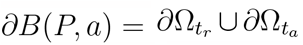. Particles diffuse freely in and out of the boundary subsection 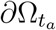, of size 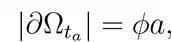 where *ϕ* is angle of the free boundary.
3. **Reflecting boundary condition on**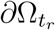 **and absorbing on** 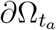 . Particles hitting the boundary subsection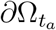 are permanently absorbed.

In the third case, to compensate for the unrealistic absorbing boundary condition, we need to accounte for absorbed particles that can also escape the target domain at a rate 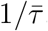, given by the narrow escape formula [23] on the disk

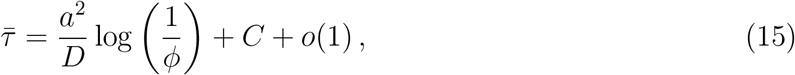

where *C* is a known constant [23]. Thus we can derive an equation for the total mass *S*_target_(*t*) inside the target region *B*(*P, a*) by substracting a loss term, that is proportional to the overall number of particles, from the influx due to the absorbing boundary condition. This balance equation is given by

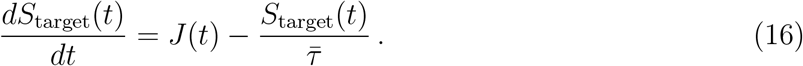

where *J* (*t*) is the time-dependent flux *J* (*t*) through the narrow window,

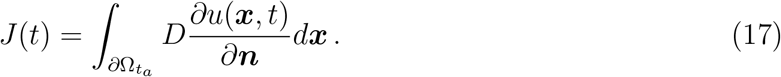

The solution for the total mass (case 3) within the target is given by

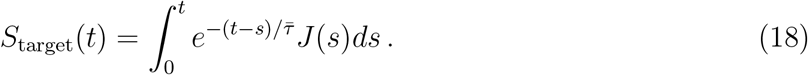

The COMSOL software allows exploring how the source-target located at a distance *L*, as well as the target permeability, controlled by the angle *ϕ* of free portion of boundary, affects the responses for the three different cases (Fig. 3). Lower responses are recorded as the source-target distance increases (case 1, Fig. 3A). Then for the second case, smaller angles *ϕ* lead to smoother target responses: it is harder for particles to penetrate the target region, but once inside they stay longer on average (Fig. 3B). Stepper response curves are also found for small angles *ϕ* when the target opening is fully absorbing, with the exit modeled using classical narrow escape theory (case 3, Fig. 3C).

**Figure 3:**
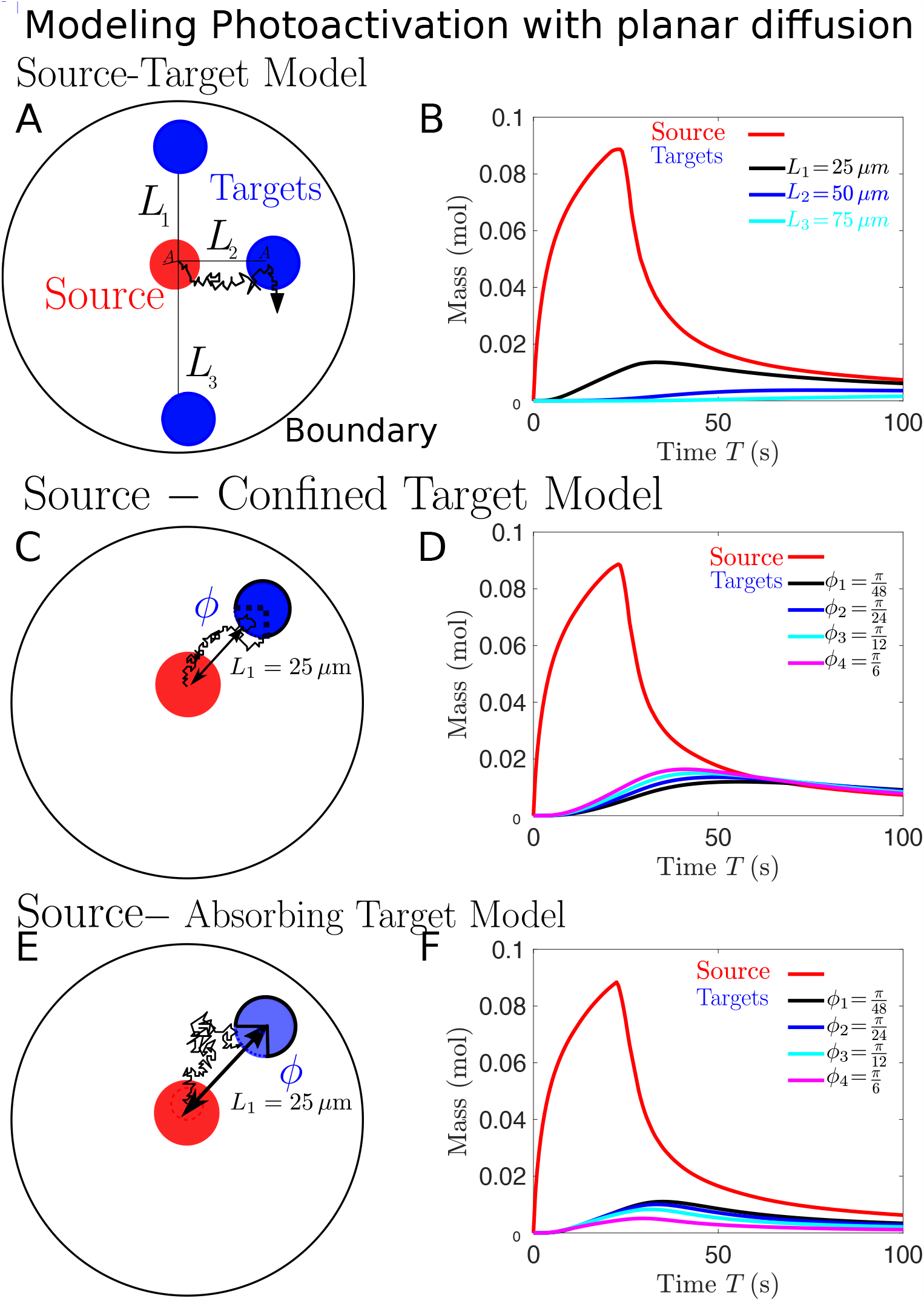
Effect of target organization and permeability on source-target responses. **A**. No further conditions are imposed on the target boundary *∂B*(*O, a*). The source is simulated (red curve) and compared to the target responses (black, dark and light blue curves) for three distances *L* = 25, 50 and 75 *µm*. **B**. Particles diffuse freely in and out of the small boundary subsection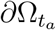 of size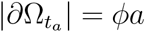, where *ϕ* is the free boundary angle, while the large remaining boundary portion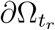 is reflecting. **C**. Particles hitting the boundary subsection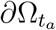 are absorbed and reinjected in the bulk at a rate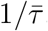, with 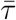 given by Eq. 15. Parameter values are *D* = 10 *µ*m^2^*/*s, *R* = 100 *µ*m, *a* = 5 *µ*m, *c*_0_ = 1 mol, *t*_*s*_ = 25 s, *γ*_*s*_ = 1 s and *γ* = 1 *µ*m.

**Figure 4:**
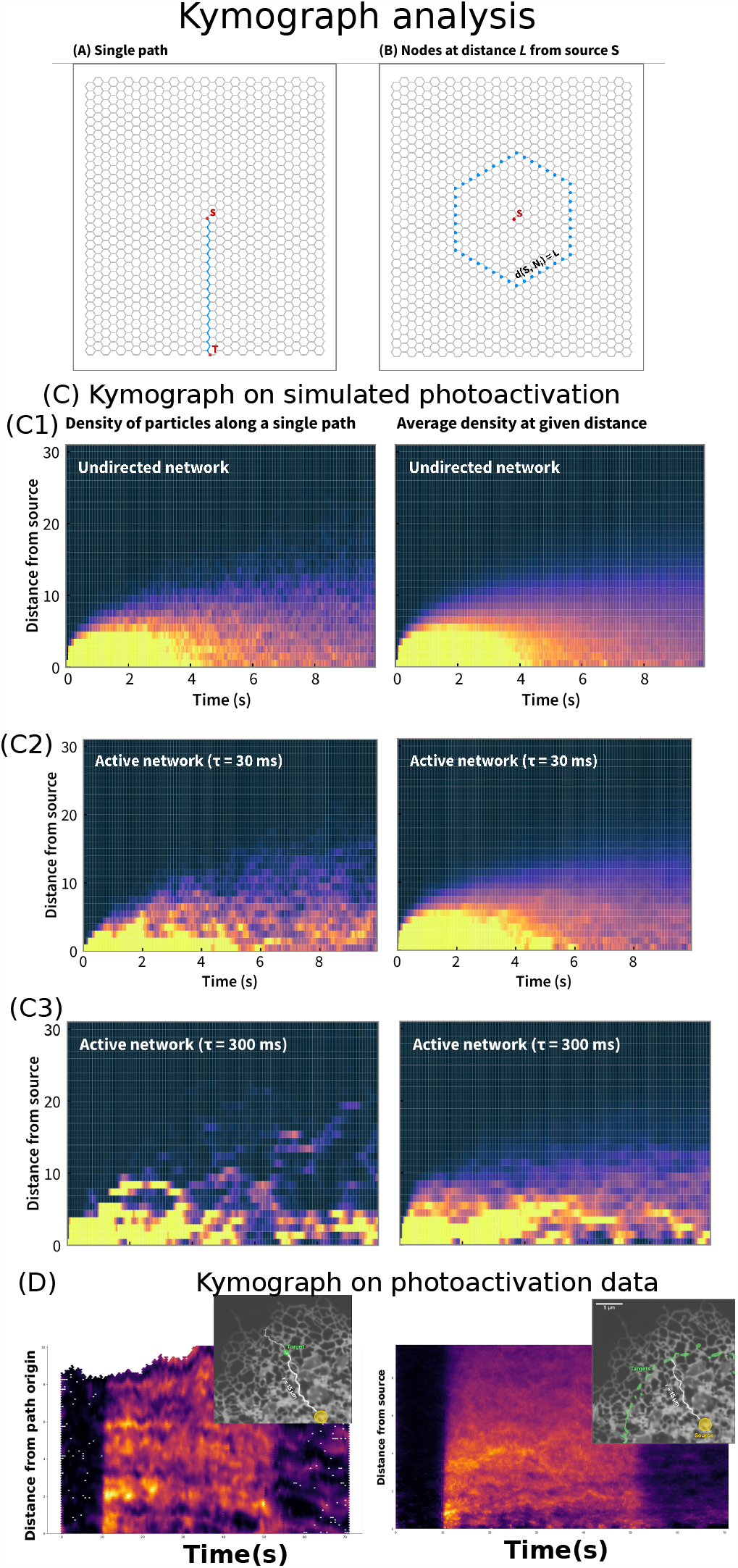
Kymograph Analysis of Photoactivation. **A-B**. Representation of the distance computed along a path or by averaging over an equidistance from the center in a tridiagonal network. **C**. Time-distance kymograph representation of the photoactivation occurring at position *X* = (0, 0) for three different propagation modes: undirected (C1), or modulated by the switching time of the unidirectional flow with *τ* = 30 ms (C2) and *τ* = 300 ms (C3). **D**. Kymograph of photoactivation data [10].

### 3.4 Modeling Photoactivation protocole by a reaction-diffusion process at the source

We now discuss briefly an alternative model of photoactivation that accounts for the decay of activated molecules inside the source. Going back to the diffusion equation 2, a source term can be added restricted to the d k of radius *R*:

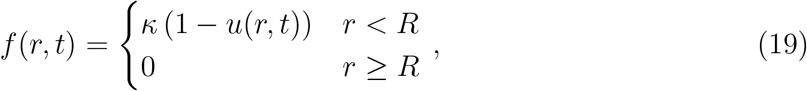

describing the activation of inactive particles by the laser beam at a rate *κ*. The dynamics of active molecules occurring in a two-dimensional plane is thus modeled by the reaction-diffusion equation:

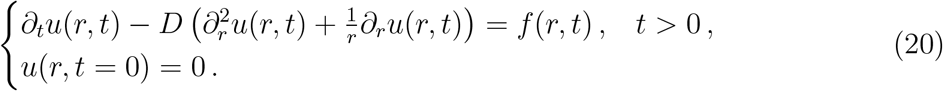

The equation is solved in two distinct regions, for *r < R* and *r < R* with solutions *u*_*r<R*_(*R, t*) and *u*_*r>R*_(*R, t*) respectively (see appendix 7.1.1). For short-time 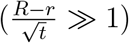, we find (see also appendix 7.1),

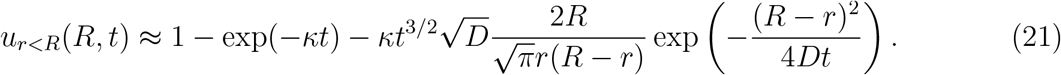

Thus to leading order, the fluorescence decay at the source can be used to estimate the rate *κ*. Outside the source, the fluorescence decay is characterized by

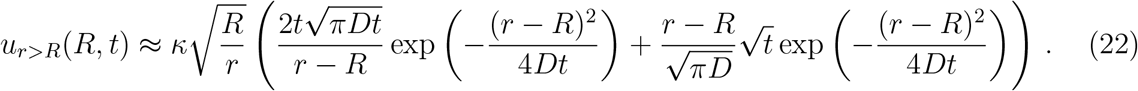

This expression shows that the decay away from the source could be used to extract the diffusion coefficient once the rate *κ* has been determined (see also appendix 7.2).

## 4 Kymograph construction of Photoactivation

We describe in this section the kymograph approach which is a time-position representation of the fluoresence. A kymograph was originally a device that draws the spatial position of a tracking object over time in which a spatial axis represents time. This time-space representation is used to study the velocity over time (continuous and jump moments) of a tracked moving object, such as cells, organelles, molecules or molecular motor [24].

A kymograph approach can be adapted to follow the distribution of fluorescent molecules over time and space, using a time-space plot of the fluorescence (color coded by intensity), measured along one dimension [25, 26, 27]. The kymograph representation can be used to follow the spread of photoactivation. However due to the network structure, the analysis has to account for the dynamics of tubular paths through the structure and the low copy number of fluorescent molecules. The distances between the PA source and a point *P* is computed from the segmented ER structure. We can also average the fluorescence extracted at subdomains located at given distance. This average kymograph summarizes the time-evolution of fluorescence versus the distance from the PA source (Fig. 4A).

This representation can be tested on simulated data [28], revealing the differences between classical diffusion and active transport (Fig. 4A-D). In particular packet transport is shown by the heterogeneous concentration distribution. In photoactivation data, the kymograph shows a concentrated signal close to the PA source and a gradual spreading through the ER, moving towards distant regions, as photoactivation increases. The time evolution of the fluorescence can be fitted by a diffusion model, ⟨*r*^2^*u*(*r, t*)⟩_*r*_ *∼*4*Dt* [10]. To conclude, the kymograph allows to quantify the spread of fluorescence across a tubular path, thus allowing to detect possible deviations from diffusion.

## 5 Modeling active versus passive transport on graph to reconstruct the photoactivation response

We discuss in this section how network modeling can be used to simulate photoactivation and trafficking inside an ER. Indeed, another possibility to model photoactivation is to use a network (see also appendix 7.3) of nodes connected by edges representing tubules (Fig. 5) [29, 30]. The degree represents the number of connections per node of the graph and thus accounts for its topology. Reconstructing a network from Structure illumination microscopy images leads to a homogeneous number of connection of degree *d* = 3. It is also possible to use a homogeneous hexagonal lattice (Fig. 5), with various conditions at the periphery nodes: molecules can be reflected, absorbed or periodic conditions can be imposed.

**Figure 5:**
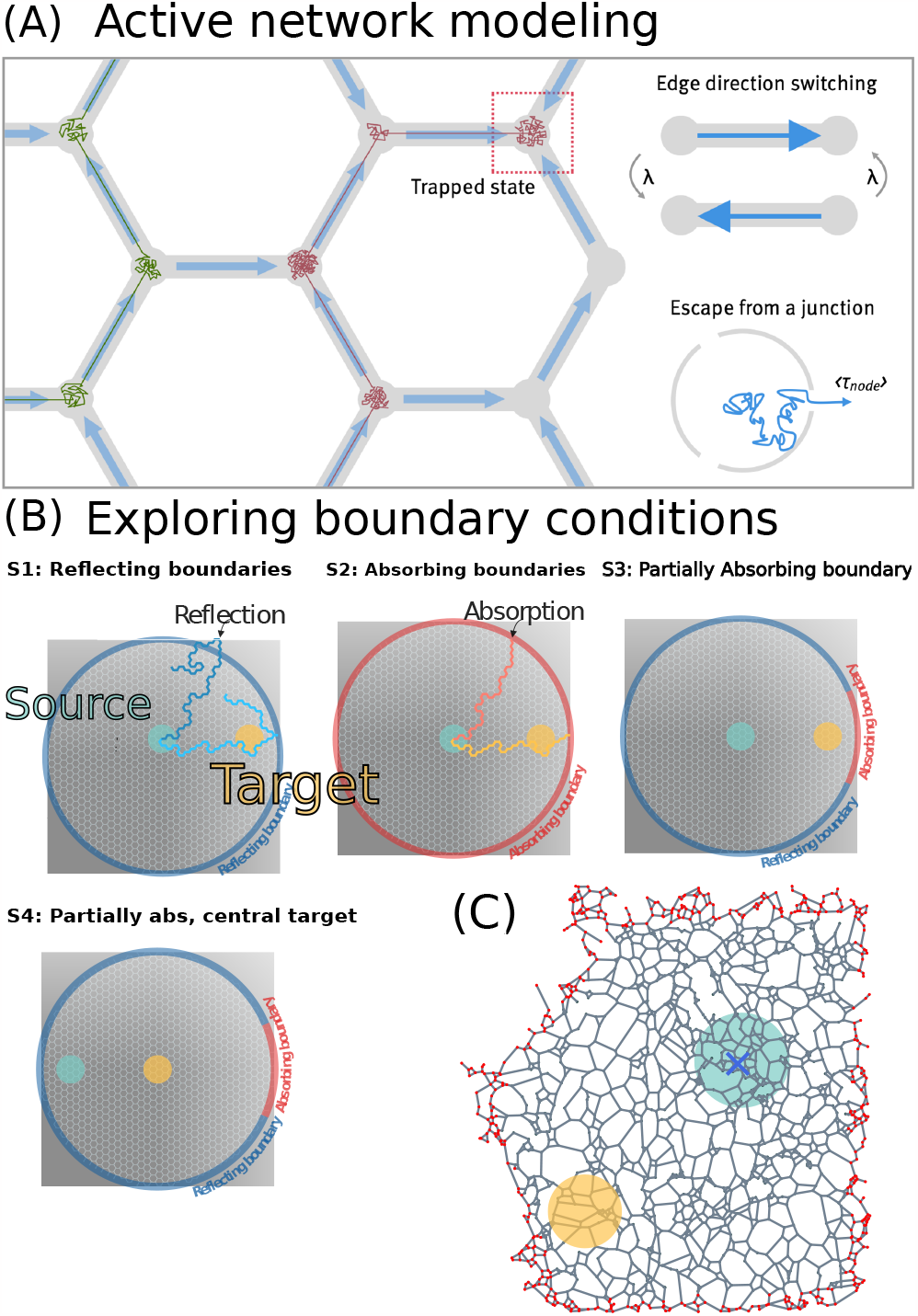
ER network modeling and conditions at boundary nodes. **A**. Active network modeling consisting of alternating flows in tubules occurring at a rate *λ*. Escape from a junction occurs with a probability which is the reciprocal of the number of connected nodes. **B**. Examples of source-target representation in an hexagonal lattice network with various boundary conditions imposed at a disk of radius R. Different boundary conditions modulate the time course of the simulated PA, allowing to explore the optimal configuration for a give PA dynamics. **C**. Reconstruction of a partial ER network from SIM (Structure illumination microscopy), where any boundary conditions can be given at external nodes (red), adapted from [28].

A network is described by a graph *G* = (*V, E*) of vertices and edges. The transition probability between nodes is described by the matrix *M* = [*m*_*ij*_]_*i,j∈V*_

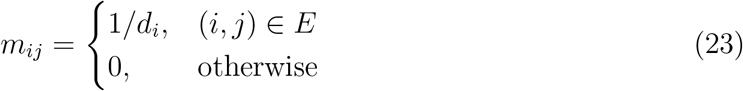

The time evolution of a transition process describes the number of molecules occupying a node at time t. In a discrete time representation, we have

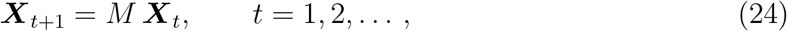

and consequently the time evolution depends on the initial distribution

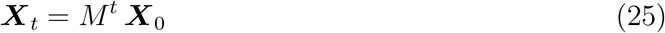

where ***X***_0_ is the initial state at time 0.

### 5.1 Diffusion-network models

Diffusion in a network consists in splitting at each node the number of particles between the number of connected nodes at either random time or synchronously in case of a simulation with a clock time ∆*t*. The time in each tubule is zero (infinite velocity). For a rate *λ* and a size *l*, the effective diffusion coefficient is *D* = *λl*^2^. Diffusion is characterized by a uniform spread of molecules until steady-state where the final density is uniform 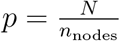, where N is the initial number of particles and *n*_nodes_ the number of nodes [28].

### 5.2 Active-network model

In an active-network model, the direction of the flow in each tubule alternates: edges have a directionality that changes in time according to a Poissonian rate. In an active-network model, a particle moves from one node to another by selecting a random incident edge from those which are outwards directed and jumps through it. When all edges are inward directed (capture state), particles at the nodes wait a random time and then retry to escape until one of the edge becomes outward (Fig. 5). Thus stochastic motion on an active network is characterized by two parameters: a tubule switching rate *λ* for tubular flow to change directions and waiting rate *λ* for particles to move to the chosen neighboring node.

### 5.3 Deconvolution of the injection rate

A first step for simulating PA responses is to reproduce the fluorescence dynamics or the number of molecules at the source. To simulate the PA process, we need to convert the measured photoconverted molecular dynamics into an injection rate. Concentration is not an injection rate due to molecules entering and exiting the source region. The injection rate *r*(*t*) should be found from the measured concentration c(t). Thus this step relies on finding the injection rate or the number of photoactivated particles. When particles are released gradually over time at a single source node *s*, then we need to replace the initial vector ***X***_0_ such that *x*_0,*i*=*s*_ = 1 and *x*_0,*i*≠*s*_ = 0 by a vector accounting for the injection rate constants. For an injection rate *µ*_*t*_ we obtain a classical convolution for the distribution of particles at time *t* as

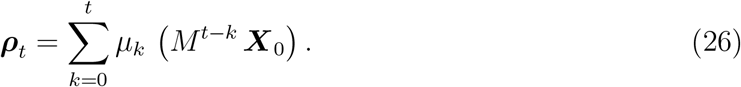

When the density *ϕ*_*t*_ of particles is known in a ROI and the injection node, the rate *µ*_*t*_ at which particles were injected can be deconvolve from eq. 26 using the vector ***π*** for obtaining the total density in the ROI.

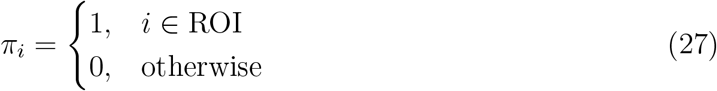

Thus,

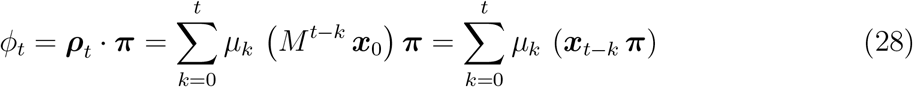

Using the first time *T* for a particle to reach the ROI from the source node, we have ***x***_*t*_ ***π*** = 0 for any *t < T*, leading to

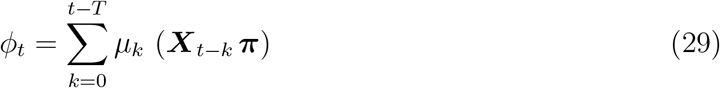

and we can recover the coefficients *µ*_*t*_ recursively, starting from 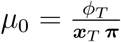 using

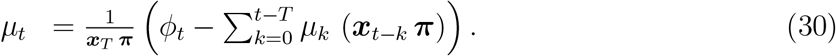

### 5.4 Simulating PA in hexagonal or reconstructed network

The flow of material is often only available in a small portion of a graph. To model photoactivation, the first step is to choose a network (hexagonal or empirical, Fig. 6A1-B1-C1). First Passage time analysis reveal how geometry can influence trafficking as shown in Fig. 6A2/A3-B2/B3-C2/C3, where simulation parameters are summarized in tables 1-2.

**Table 1:** Network structure characteristics.

| Structure | Nodes | Edges | Radius | Degree (avg) |
| --- | --- | --- | --- | --- |
| Hexagonal lattice | 2046 | 3006 | 47 | 2.94 |
| Periodic hex. lattice | 1984 | 2976 | 47 | 3 |
| Reconstructed ER | 1920 | 2787 | 41 | 2.90 |

**Table 2:** Simulation parameters.

| Param | Value | Distribution |
| --- | --- | --- |
| $\tau_{edge}$ | 0 | Constant |
| $\tau_{node}$ | 100ms | Exponential |
| $\tau_{switch}$ | 30ms | Exponential |

**Figure 6:**
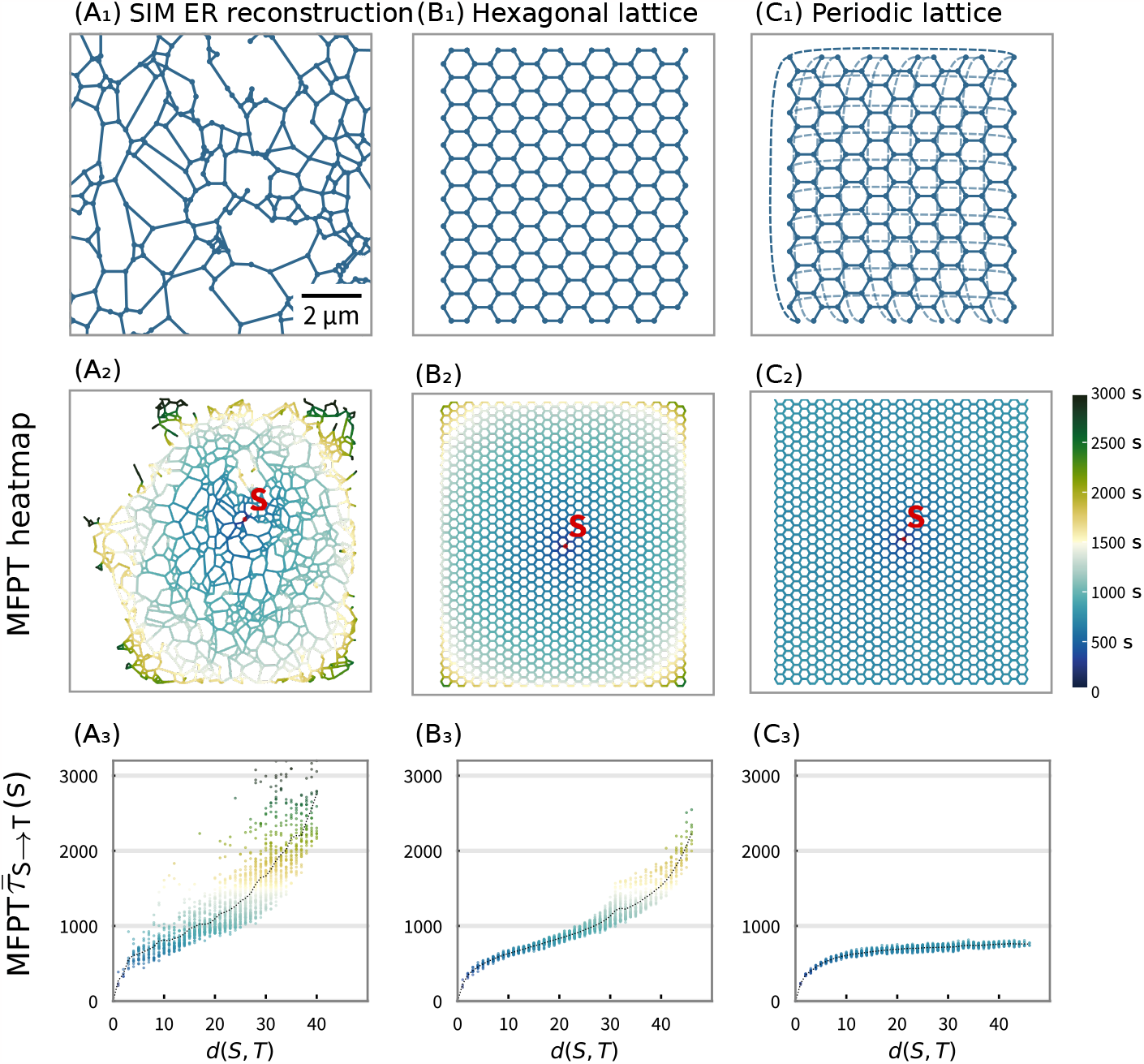
Stochastic simulations of arrival times in three graphs. **A** Three examples of network reconstruction: *A*_1_ from Structure illumination microscopy data, *B*_1_ Hexagonal lattice with reflecting boundary on the edge nodes *C*_1_ Hexagonal lattice with periodic boundary. **B** Probability maps of the first arrival time depending on the distance from the source *S* for the three reconstructed graphs. **C** Distribution of the Mean First Passage Times (MFPT) vs distances. Parameters are summarized in tables 1-2.

Once a model is selected, which can either be diffusion-based or with an active flow controlling directionality [28], the PA in the source region can be used to deconvolve the injection rate and thus to obtain a first photoactivation approximation. Once the source is well calibrated, the model parameters should be adjusted such that the simulations and the PA responses at the source match. Diffusion on a graph is simulated as a jump process between nodes: with a mean distance between neighboring nodes of around *L*_*ER*_ = 1 *µ*m [9, 18] with an exponential waiting time with a mean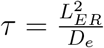 .

Two examples are shown in Fig. 7, with such an analysis allowing to recover the diffusion coefficient. Alternatively for an active flow model, the two parameters to adjust are the *µ* and *λ* switching rates. Another free parameter of the simulation is the flux condition at nodes located on the boundary of the simulation region. These boundary conditions can influence both the rising and the decay phases. Several scenario are possible: the flux cannot (resp. can) return to the domain when reaching a boundary point, described as purely absorbing (resp. reflecting) points. In an intermediate case, a fraction *P* of the flow is reflected and the rest 1*− P* is absorbed.

**Figure 7:**
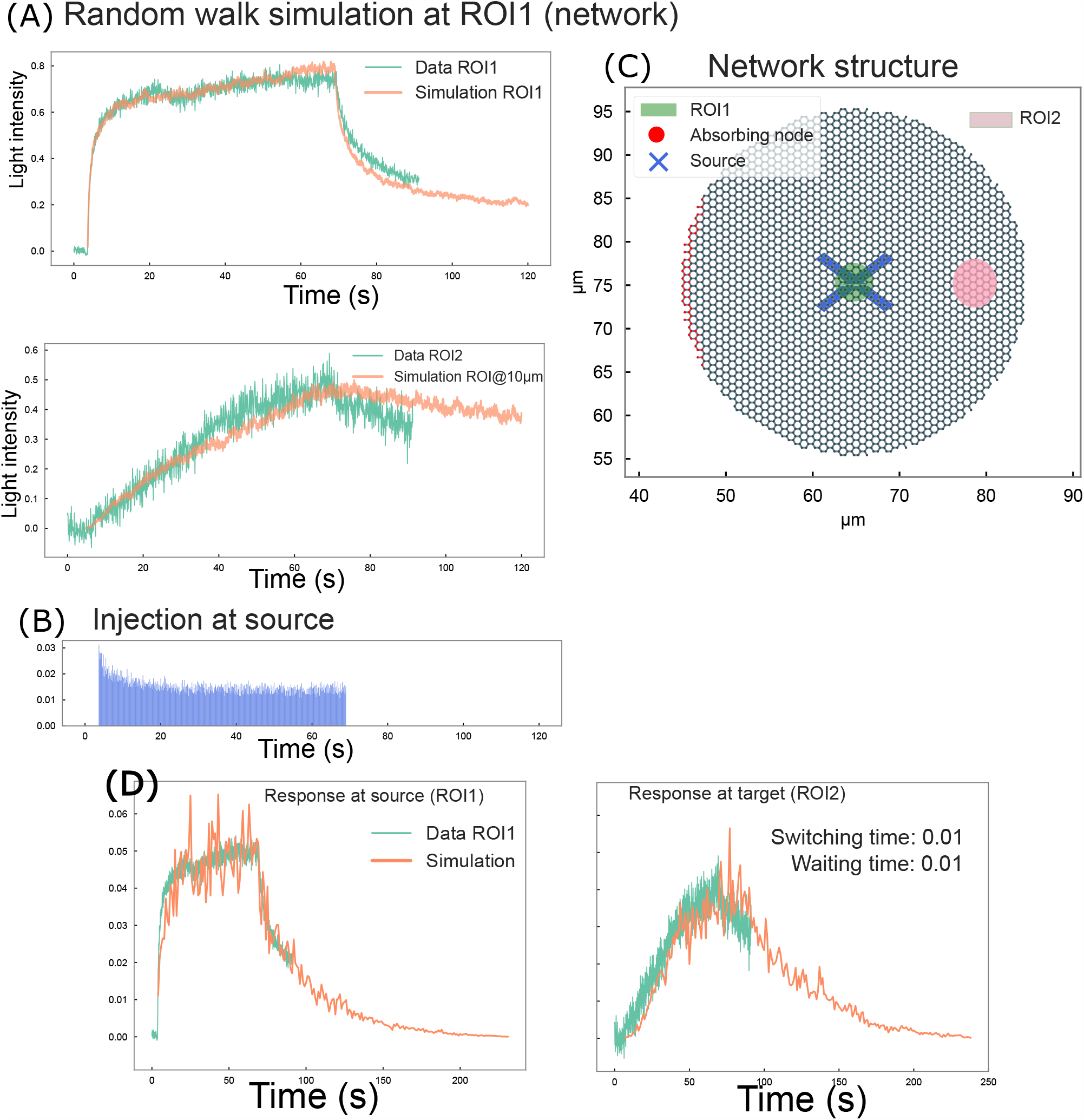
Stochastic simulations of photoactivation responses. **A-B**. Photoactivation at the source (ROI1) and the target (RO2)in green versus random walk simulations for an injection rate described by eq. 30. **C**. Brownian simulations in a hexagonal network, where the source (green) is at the center and the target is ROI2 (purple). The boundary conditions on the nodes are either reflecting (black) or absorbing (black). **D**. Source-target responses within an active flow network where similar switching and waiting times 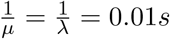.

Transients during the initial Activation-Saturation phases are not sensitive to the boundary condition imposed on the partially simulated network, because it depends mostly on the local properties of the network and laser conversion rate. Interestingly, the boundary condition influences drastically the recovery phase and the dynamics at the target (Fig. 7). It remains difficult to recapitulate the PA dynamics with the present diffusing network simulations. Another example concerns an active network with a switching time of 0.01 *s* and waiting time of 0.01 *s*, when the distance between the source and the target is *d* = 23 *µ*m, with a target radius of 2.2 *µ*m. Results are shown in Fig. 7.

## 6 Discussion

We presented here computational approaches based on solving diffusion equations, computing Mean Square Displacement (MSD) or Kymograph analyses [28, 10], in order to analyze the first phases of photoactivation. These approaches are used to characterize molecular transport in the endoplasmic reticulum. They account for fluorescence data at a source and a target sites. Biophysical based models rely on classical and anomalous diffusion. We presented here how these methods are used to extract various parameters, such as the anomalous exponent, the diffusion coefficient and the diffusion map.

It is possible to adapt these methods to study the decay phase of the PA, of which it would be interesting to extract the associated biophysical parameters. In particular using stochastic simulations in a network allows exploring different boundary conditions that should be imposed at each peripheral node to recover the PA decay: this could be a mix of absorbing or partial absorbing conditions with reflecting. These conditions affect the decay rate at short time, and thus could reveal hidden control mechanisms of local transport.

Previous stochastic modelings [28] explored the possibility that molecules in the ER lumen could traffick together in packets. This transport is in contrast with classical diffusion laws, characterized by molecular dispersion. So far, local transiently persistent fluorescence was replicated by solving the diffusion equation in a heterogeneous network, accounting for local differences in molecular accumulation in nodes [10]. This approach can rely on an image processing procedure to remove noise and to account for the continuously changing ER morphology during lumen trafficking, an approach that will be discussed somewhere else. Once key parameters are extracted, computations and numerical simulations can be used to estimate arrival rates to any regions of interest and predict the accumulation of molecules at exit sites. Another application of the computational methods presented here would be to develop simulations of misfolded protein trafficking. A possible model consists in diffusion-aggregation, where the diffusion coefficient could depend on the size of aggregates [12]. It would also be possible to apply the present method to quantify the effect of ATP depletion on PA to explore the role of possible active components.

## 6.1 Acknowledgement

We will to Thank Chris Obara for critical feedback on this manuscript. This project has received funding from the European Research Council (ERC) to D.H. under the European Union’s Horizon 2020 research and innovation programme (grant agreement No 882673) and ARC and ANR NEUC 00001 and ANR AstroXcite and ARC PDF12020020001505 to F.P.L.

## 7 Appendix

### 7.1 Short-time behavior of reaction-diffusion approximation to photo-activation

Eq. 20 is solved by separately considering the two cases *r < R* and *r≥ R*. For *r≥R*, the homogeneous partial differential equation is

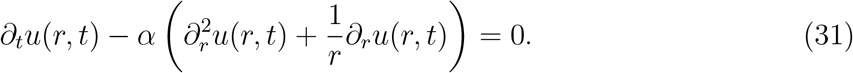

where the diffusion coefficient is now denoted by *α*. By then taking the Laplace transform, we obtain the modified Bessel equation

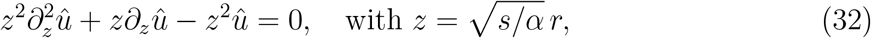

which has a general solution

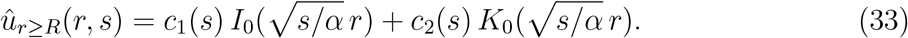

In the case *r < R*, we need to solve the nonhomogeneous equation:

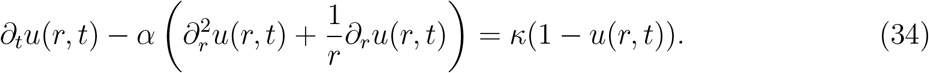

By taking the Laplace transform, we obtain

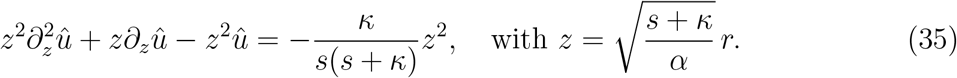

The general solution of eq. 35 is the sum of the solutions of the corresponding homogeneous equation and a particular solution *ψ* _part_. The corresponding homogeneous equation is a modified Bessel equation with solutions *I*_0_(*z*) and *K*_0_(*z*). The particular solution is constant in *r* and 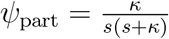. The solution for the case *r < R* is then:

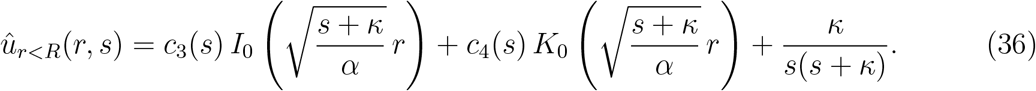

Putting the two solutions together, the solution of eq. 20 in the Laplace domain is

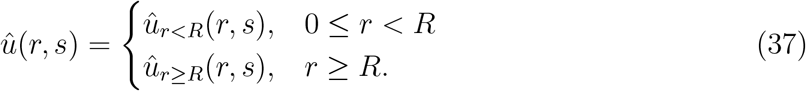

The coefficients *c*_1_, *c*_2_, *c*_3_, *c*_4_ are determined by we constraints on the solution: it has to be bounded in *r*, leading to *c*_4_ = 0 prevent a diverging behavior at *r* = 0. Similarly, *c*_1_ = 0 to prevent the solution to diverge for *r→ ∞* . The remaining coefficients are found by using the continuity of the solution and its first derivative at *r* = *R*:

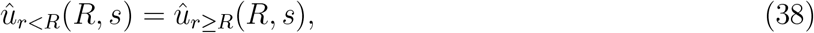

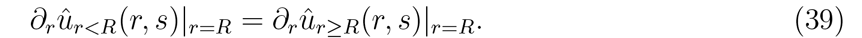

Finally, we get

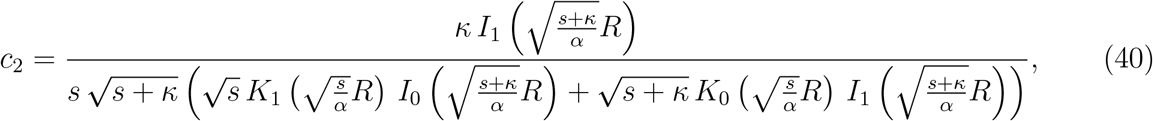

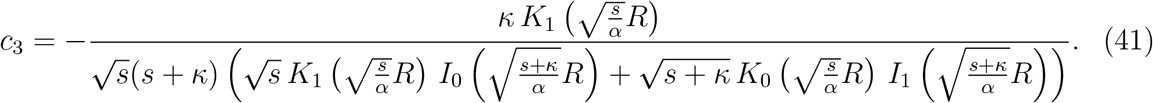

### 7.1.1 Short-time asymptotics of reaction-diffusion equation

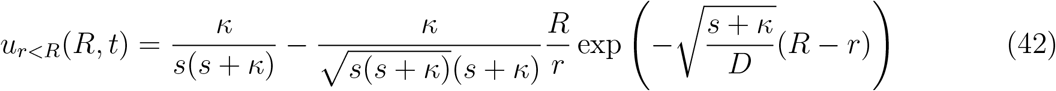

and

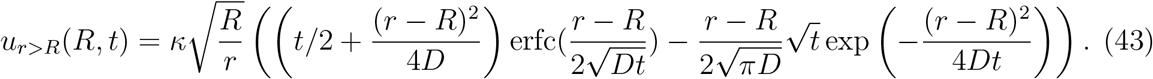

### 7.2 Short-time behavior of diffusion approximation to photo-activation

When photoactivation stops at the source, we can use as an approximation the solution of a diffusion equation where a concentration *C*_1_ is maintained constant on the disk of radius *r* = *a*. The radial solution is well-known [20] (p. 87) to be

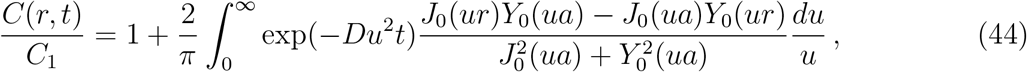

where *J*_0_, *Y*_0_ are the bessel functions. For short-time, the decay at the source is approximated by

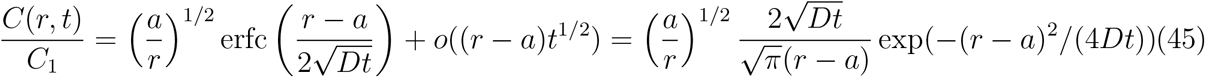

This expression could be used to fit the diffusion coefficient for a dense ER network, so that the possible ER does not impact the two-dimensional planar approximation.

### 7.3 Modeling diffusion and active transport in a graph

We recall here some general theory about transport in a network. The network is represented by the matrix *A*_*pq*_ accounting for the topology (specific nodes to be connected), where *A*_*pq*_ = 1 if the nodes *p* and *q* are connected and *A*_*pq*_ = 0 otherwise. The transition probability density function *P*_*p*_(*t* + ∆*t*) for a particle ending at time *t* + ∆*t* at node *p* is given by the sum of the transition rates *λ*_*p→q*_(*t*) from *p* to *q* at time t multiplied by the probability that a particle is at node q at time t:

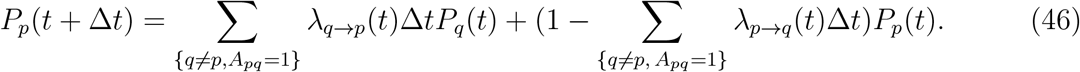

The transition rates are

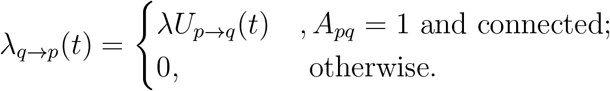

and the random Poissonian variable satisfies

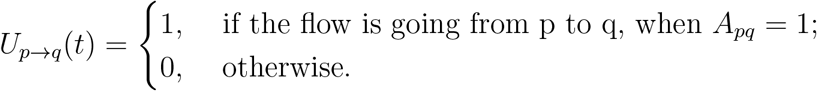

The pdf of *U*_*p→q*_(*t*) reflects the switching flow between two connected nodes. It is Poissonian with rate *λ*_*sw*_, so that the Markov chain for the pdf *P*_*p→q*_ is independent of the node and tubule and is given by *P*_*p→q*_ + *P*_*q→p*_ = 1 and

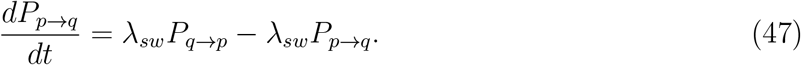

If a particle is ready to escape from a node, but all tubules are characterized by converging flow direction, the particle will have to wait that the flow in one of the tubule changes direction in order to be expelled from the node to the next one. Thus the escape rate *λ* is such

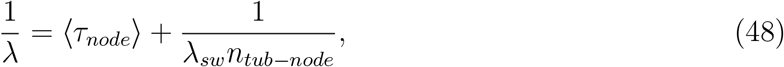

where ⟨*τ*_*node*_⟩ is the mean residence time in a node, *n*_*tub−node*_ is the number of tubules per node (equal to 3 here). When the switching time is much larger than *(τ*_*node*_), this condition models extra-long confinement. In the opposite regime, where the 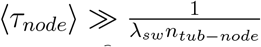, the process is equivalent to a random walk on a graph. In that case, the mean first passage time from a node to a given one can be computed asymptotically for an hexagonal lattice [28].

